# Circulating APOH promotes aortic dissection by activating the vascular smooth muscle cell NR5A1–PPARγ pathway

**DOI:** 10.64898/2026.07.16.739043

**Authors:** Likang Ma, Lei Jin, Juncheng Liu, Jiakang Li, Maolin Liu, Liangwan Chen, Zhihuang Qiu

**Affiliations:** Department of Cardiovascular Surgery, Fujian Medical University Union Hospital, Fuzhou 350001, China; Fujian Medical University Heart Center, Fuzhou 350001, China; NHC Key Laboratory of Etiological Epidemiology of Chronic Diseases with High Incidence in Fujian-Taiwan Area (Co-construction), Fujian Medical University, Fuzhou 350122, China; The Key Laboratory of Fujian Province Universities on Ion Channel and Signal Transduction in Cardiovascular Diseases, Department of Physiology and Pathophysiology, The School of Basic Medical Sciences, Fujian Medical University, Fuzhou 350001, China; Department of Outpatient, Fujian Medical University Union Hospital, Fuzhou 350001, China

**Keywords:** Aortic dissection, Apolipoprotein H, Vascular smooth muscle cell phenotypic switching, NR5A1–PPARγ axis, Inflammation

## Abstract

**Introductions:** Aortic dissection (AD) is a life-threatening vascular disease with limited therapeutic targets. Apolipoprotein H (APOH), a circulating glycoprotein implicated in lipid metabolism, has not been studied in AD.

**Methods:** Plasma APOH levels and aortic deposition were examined in AD patients. A β-aminopropionitrile (BAPN) and angiotensin II (Ang-II)-induced mouse AD model with AAV-mediated Apoh knockdown was used to evaluate survival, aortic dilation, and extracellular matrix remodeling. Transcriptomic profiling, chromatin immunoprecipitation, and gene silencing in human aortic vascular smooth muscle cells (HAVSMC) were performed to dissect the mechanism. PPARγ agonist rescue was conducted *in vivo*.

**Results:** APOH was elevated in plasma and deposited in AD aortas. Apoh knockdown improved survival, reduced AD incidence and ascending aortic dilation, and attenuated elastic fiber disruption and collagen deposition. Transcriptomics revealed enrichment of the PPAR pathway. APOH promoted HAVSMC phenotypic switching from a contractile to a synthetic state, decreasing ACTA2/TAGLN and increasing OPN/MMP9. Mechanistically, APOH upregulated NR5A1, which directly bound the PPARγ promoter to enhance PPARγ and FABP4 expression. Silencing NR5A1 or PPARγ reversed APOH-induced phenotypic switching and inflammation. *In vivo*, PPARγ agonist diminished the protective effects of Apoh silencing.

**Conclusion:** APOH promotes AD progression through the NR5A1–PPARγ axis, driving vascular smooth muscle cell phenotypic switching and inflammation, and represents a potential therapeutic target.

**Graphical Abstract:** 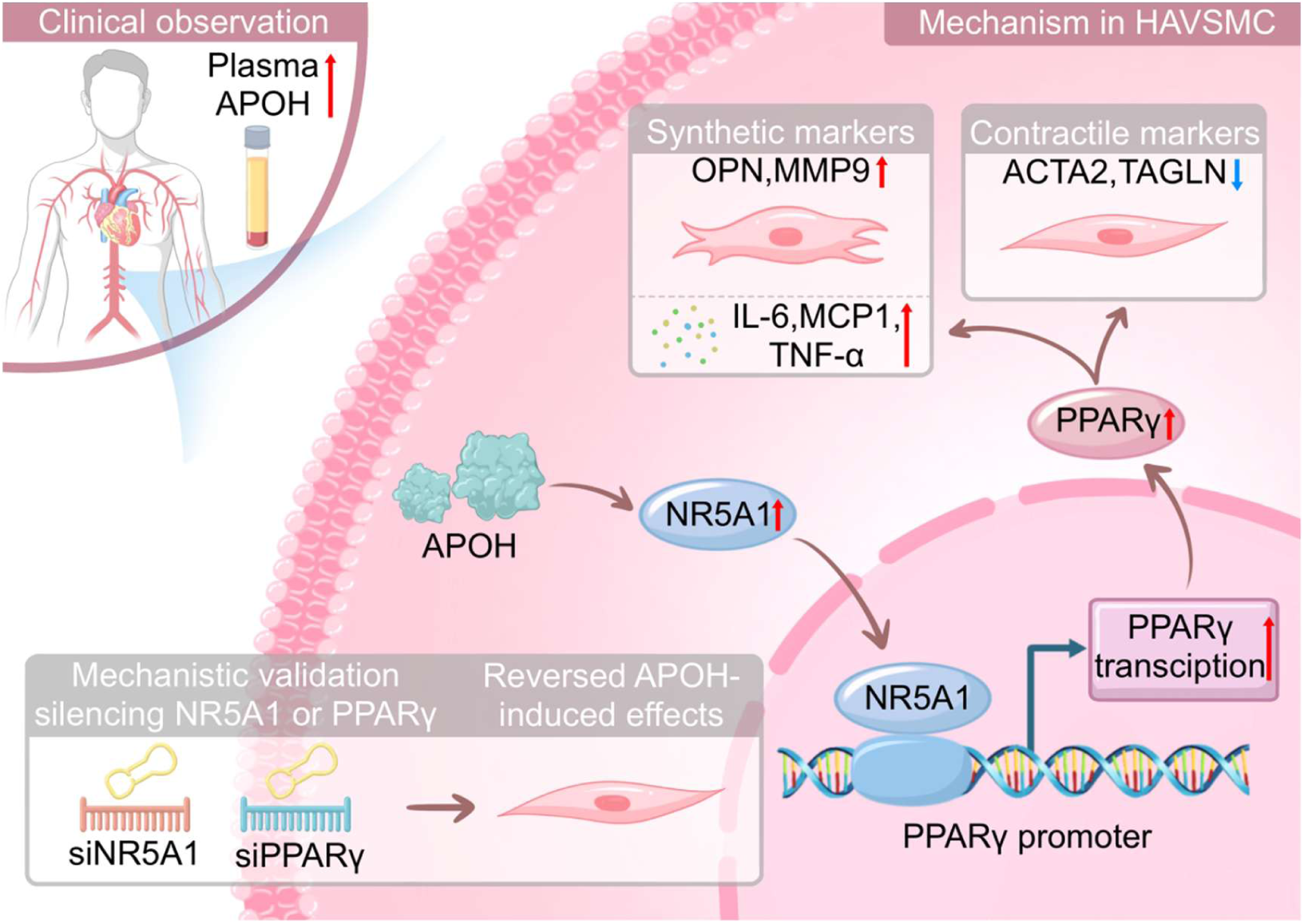

**Circulating APOH promotes aortic dissection through the NR5A1–PPARγ axis in human aortic vascular smooth muscle cells.**

Clinical observations showed that plasma APOH levels were elevated in patients with aortic dissection. Circulating APOH acts on human aortic vascular smooth muscle cells (HAVSMC) and upregulates NR5A1, which binds to the PPARG promoter and enhances PPARγ transcription. Activation of the NR5A1–PPARγ signaling axis promotes the phenotypic transition of HAVSMCs from a contractile phenotype to a synthetic phenotype, as indicated by decreased ACTA2 and TAGLN expression and increased OPN and MMP9 expression, accompanied by enhanced production of the inflammatory mediators IL-6, MCP-1, and TNF-α. Silencing NR5A1 or PPARγ reverses APOH-induced phenotypic switching and inflammatory responses, supporting the critical role of the NR5A1–PPARγ axis in APOH-mediated vascular injury.

## INTRODUCTION

Aortic dissection (AD) poses a serious threat to human health with an acute onset, high mortality rate, and no reliable molecular prevention or treatment targets^1,2^. Degradation of the middle layer of the aorta and imbalance in extracellular matrix (ECM) synthesis and degradation are the main pathological changes in AD^1,3^. Collagen and elastin are the main components of the aortic wall, accounting for 50% of the normal arterial dry weight. They play an important role in maintaining the integrity of the aorta and withstanding the stress of blood flow on the wall^3^. Numerous studies have shown that the arterial wall of patients with AD becomes thinner, with elastic protein fragments visible in the middle layer, and an imbalance in the elastin-to-collagen ratio^3^. This pathological change increases the aortic compliance, which in turn can exacerbate mid-layer degeneration, leading to further elastin fiber defects and ultimately degradation of the basement membrane, thereby resulting in aortic dissection^1–4^.

Human aortic vascular smooth muscle cell (HAVSMC) phenotypic switching plays a critical role in maintaining the structural homeostasis of the aortic wall. Under physiological conditions, HAVSMC exhibit a contractile phenotype, which is responsible for maintaining vascular tone and ECM homeostasis. Upon pathological stimulation, HAVSMC undergo phenotypic switching toward a synthetic phenotype, characterized by downregulation of contractile marker genes and increased secretion of matrix-degrading enzymes, thereby exacerbating ECM degradation and medial degeneration^5^. Among these enzymes, matrix metalloproteinase 9 (MMP9) serves as a key effector molecule during HAVSMC phenotypic switching and participates in the regulation of ECM degradation^6^. In patients with AD, MMP9 expression is significantly elevated in the aortic wall and serum, and its increased activity contributes to enhanced aortic compliance, aggravated collagen degradation, and aortic wall thinning, thereby promoting AD progression^6^.

Recently, glucose and lipid metabolism have been implicated in AD pathogenesis. Succinate levels are elevated in the plasma of AD patients, and lowering succinate alleviates disease progression^7^. LHDA, Anxa1, and high-density lipoprotein (HDL) regulate HAVSMC phenotypic switching via modulating glucose and lipid metabolism, thereby contributing to AD^8,9^. These findings suggest that metabolic abnormalities influence AD development, at least in part, through HAVSMC phenotype regulation. Based on the above background, the present study performed sequencing analysis of plasma proteins and RNA from patients with AD and healthy controls, and identified significantly upregulated expression of apolipoprotein H (APOH). APOH, also known as β2-glycoprotein I, is a component of circulating plasma lipoproteins, primarily synthesized in the liver, and is involved in triglyceride metabolism and the regulation of lipid metabolism^10^. However, no studies to date have reported on the role of APOH in the pathogenesis of AD. This study aims to investigate the potential regulatory mechanism of APOH in the development and progression of AD.

## METHODS

### Clinical samples collection

This study included patients with aortic dissection (AD) diagnosed using computed tomography (CT) imaging in the Cardiac Surgery Department of Fujian Medical University Union Hospital between January 2024 and June 2025. Written informed consent was obtained from all participants for the use of human blood and tissue samples. AD tissue samples were obtained from the thoracic aorta of AD patients undergoing surgery; specifically, they were taken from the dissected aortic wall at the site of the intimal tear or the most severely affected segment, and after resection all tissues were transferred to liquid nitrogen or RNAlater within 30 minutes for subsequent RNA extraction, with some tissues also stored in 4% paraformaldehyde for pathological staining. Normal aortic tissues were obtained from heart transplant recipients without vascular disease; although these recipients may have primary cardiomyopathy, we only included those with no pre-existing aortic pathology, atherosclerosis, or macrovascular disease, as confirmed by preoperative imaging (echocardiography and/or CT) and intraoperative inspection. The specific exclusion criteria for these normal tissue controls were: any abnormality of the aorta (e.g., dilation, aneurysm, dissection, coarctation) on imaging, history of hypertension, or dyslipidemia (elevated total cholesterol, LDL, or triglycerides; low HDL). Plasma samples from AD patients and the healthy volunteers were collected. The healthy volunteers were recruited during routine health check-ups in the same period, had no history of cardiovascular disease, diabetes, liver disease, or inflammatory disease, and were age- and sex-matched to the AD group. The research protocol was approved by the Ethics Committee of Fujian Medical University Union Hospital [No.2023-96]. Individual clinical characteristics of patients are provided in supplemental Table S1.

### Animal models

Animal experiments were approved by the Animal Ethics Committee of Fujian Medical University [IACUC FJMU 2023-Y-0269] and all procedures complied with the ARRIVE guidelines and relevant regulations. C57BL/6 mice were obtained from Beijing HFK Bioscience Co., Ltd. (Beijing, China). AAV9 carrying shRNA targeting mouse *Apoh* (AAV^sh-Apoh^) or scrambled control (AAV^sh-NC^) was provided by Applied Biological Materials (Shanghai, China). Mice were injected via the tail vein with 1×10¹¹ viral genomes per mouse in 50 μL PBS at 2 weeks of age and then allowed to grow for 2 weeks. For the initial animal study, mice were divided into four groups (n=15/group): Vehicle+AAV^sh-NC^, Vehicle+AAV^sh-Apoh^, BAPN+AAV^sh-NC^, and BAPN+AAV^sh-Apoh^. From 4 weeks of age, mice in BAPN-treated groups received 0.3% β-aminopropionitrile (BAPN, TCI, Japan; Cat#A0408) in drinking water for 4 weeks, followed by intraperitoneal injections of angiotensin II (Ang-II, GLPBIO, USA; Cat# GP10023) at 1.44 mg/kg three times daily (morning, noon, evening) for 3 consecutive days. Vehicle groups received normal drinking water (without BAPN) and intraperitoneal injections of PBS. This protocol avoids chronic pain associated with subcutaneous pumps, meets animal welfare requirements, and was validated by our preliminary experiments^11,12^. In the rescue experiment, all groups of mice were subjected to the same BAPN+Ang-II treatment (i.e., all developed the AD model) and received different interventions according to their group assignment (n=15/group): NC group (model only, no virus or agonist); AAV^sh-Apoh^ group (AAV^sh-Apoh^ injection via the tail vein at 2 weeks of age, otherwise the same as the model); AAV^sh-Apoh^+agonist^PPAR^ group (AAV^sh-Apoh^ injection at 2 weeks of age combined with daily oral gavage of rosiglitazone (10 mg/kg/day) starting from 4 weeks of age until the end of the experiment; rosiglitazone is a selective PPARγ agonist); and agonist^PPAR^ group (rosiglitazone alone at the same dose by daily oral gavage from 4 weeks of age, no virus injection). Rosiglitazone was purchased from MedChem Express (USA; Cat# HY-17386). Mice were euthanized 24h after the last intervention, and aortic tissues were collected. During the modeling period, cardiac ultrasound monitoring was performed on the mice. At the end of the experiment, mice were euthanized, and the occurrence of AD was confirmed by gross dissection. If a mouse died before the end of the experiment, the cause of death was determined immediately. The time and incidence of AD in the mice were recorded throughout the experiment. After modeling, the mouse aortas were preserved in liquid nitrogen or paraformaldehyde for subsequent analyses. Partial aortic tissues from some mice were sent to TIANGEN BIOTECH CO., LTD. (Beijing, China) for transcriptome sequencing.

### Cell culture

Primary human aortic vascular smooth muscle cells (HAVSMC, Cat# CP-H081) were purchased from Pricella Biotechnology Co., Ltd. (Wuhan, China). Cells were cultured in complete human aortic smooth muscle cell medium (Cat# CM-H081), which consists of basal medium supplemented with 5% fetal bovine serum (FBS), 1% penicillin-streptomycin solution, insulin, and smooth muscle cell growth supplement. Prior to drug treatment, cells were starved in serum-free DMEM for 4 hours. Subsequently, cells were treated with angiotensin II (Ang-II, Tocris, UK, Cat# 1158), recombinant APOH protein (MedChem Express, USA, Cat# HY-P7533) at 1 µg/ml, or respective vehicle control (culture medium). After treatment for 72 hours, cells were harvested for downstream experiments.

### siRNA transfection

HAVSMC were seeded into 6-well plates at a density of 5 × 10⁵ cells per well. On the following day, when cells reached 70%–80% confluence, siRNA transfection was performed. Prior to transfection, the culture medium was replaced with 2 mL of complete medium containing 10% fetal bovine serum. For a single well, approximately 125 μL of DMEM without antibiotics or serum was added to a centrifuge tube, followed by 100 pmol of siRNA targeting NR5A1 or PPARγ (Applied Biological Materials, Shanghai, China) with gentle mixing. Then, 4 μL of Lipofectamine 8000 (Cat# C0533-0.5 mL, Beyotime Biotechnology, Shanghai, China) was added and mixed thoroughly. The mixture was incubated at room temperature for 20 minutes, after which the transfection complex was added dropwise to the well and gently swirled. After 4 hours of transfection, the medium was replaced with fresh complete medium, and the cells were cultured for an additional 24 hours before subsequent experiments. A negative control was performed using non-targeting siRNA (si-NC) following the same procedure.

### Quantitative real-time PCR (qPCR)

Total RNA was extracted from tissues or cells using TRIzol reagent (Invitrogen, USA). Reverse transcription was performed to synthesize cDNA using a reverse transcription kit (Takara Bio, USA). Quantitative real-time PCR was carried out using TB Green Premix Ex Taq (Takara Bio, USA) on a LightCycler 480 system (Roche LC480, software release 1.5.0). The thermal cycling program was as follows: initial denaturation at 95 °C for 5 min; 40 cycles of 95 °C for 30 s, 61 °C for 30 s, and 72 °C for 30 s; followed by a melting curve analysis: 95 °C for 60 s, 55 °C for 30 s, and 95 °C for 30 s. Relative mRNA expression levels were calculated using the 2^−ΔΔCt^ method. The primer sequences used for qPCR are listed in supplemental Table S2.

### Enzyme-linked immunosorbent assay (ELISA)

APOH concentrations in serum were measured using a commercial human Apo-H SimpleStep ELISA Kit (Abcam, USA; Cat# ab274403) according to the manufacturer’s instructions. This kit employs a sandwich ELISA format with a single-wash, 90-min protocol. Briefly, 96-well plates pre-coated with anti-APOH capture antibody were incubated with serum samples for 10 min at room temperature, followed by addition of a mixture of detection antibody and horseradish peroxidase (HRP) conjugate and further incubation for 90 min at room temperature. After a single wash step, 3,3′,5,5′-tetramethylbenzidine (TMB) substrate was added and incubated for 10 min at room temperature in the dark. The reaction was terminated with stop solution, and the optical density (OD) was measured at 450 nm using a microplate reader. Sample concentrations were calculated by interpolation from a standard curve. All measurements were performed in duplicate.

### Western blot (WB)

Proteins were extracted from tissues or cells using standard procedures, mixed with SDS-DTT loading buffer, and boiled. After separation by SDS-PAGE and electrotransfer, PVDF membranes were blocked with 5% non-fat milk in TBST (Tris-buffered saline with 0.1% Tween-20). Membranes were then washed with TBST and incubated with primary antibodies overnight at 4 °C. Following primary antibody incubation, membranes were washed and incubated with appropriate secondary antibodies. Protein bands were visualized using enhanced chemiluminescence (ECL). Detailed information on all primary and secondary antibodies is provided in supplemental Table S2.

### Immunohistochemistry (IHC)

All human and mouse aortic tissues were obtained from the ascending/thoracic aorta. Each sample was sectioned into three consecutive slices, and three high-power fields were randomly selected from each slice for quantitative analysis. Tissues were embedded in paraffin, sectioned at 4 µm thickness, and baked at 60 °C. Sections were then deparaffinized, rehydrated, subjected to antigen retrieval, and blocked. Thereafter, sections were incubated with primary antibodies overnight at 4 °C. After washing with PBS, sections were incubated with appropriate secondary antibodies. Following 3,3′-diaminobenzidine (DAB) staining and nuclear counterstaining, slides were mounted, and images were captured using a panoramic MIDI scanner (3DHISTECH). Immunohistochemical staining was semi-quantitatively assessed using the H-score method^13^, calculated as: 1×(% weak) + 2×(% moderate) + 3×(% strong), ranging from 0 to 300. Detailed information on the antibodies used is provided in supplemental Table S2.

### Immunofluorescence (IF)

Tissue sections were processed as described for IHC, except that fluorescent secondary antibodies were used after overnight primary antibody incubation. Slides were then counterstained with DAPI, mounted, and scanned using the Pannoramic MIDI scanner (3DHISTECH). Details of the antibodies used are provided in supplemental Table S2.

### Histopathological staining

Hematoxylin and eosin (HE) staining: Sections were deparaffinized, rehydrated, stained with hematoxylin, differentiated, stained with eosin, dehydrated, and mounted. Elastic Van Gieson (EVG) staining: Deparaffinized and rehydrated sections were stained using a Collagen Fiber and Elastic Fiber Staining Kit (Solarbio, Beijing, China; Cat# G1597) according to the manufacturer’s protocol. Masson’s trichrome staining: After deparaffinization and rehydration, sections were stained using a commercial Masson’s trichrome staining kit (Solarbio, Beijing, China; Cat# G1340). All stained sections were scanned using a Pannoramic MIDI scanner (3DHISTECH). EVG and Masson’s trichrome staining were used to quantitatively assess the elastic fiber and collagen volume fractions, respectively. The volume fraction was calculated as the ratio of elastic fiber or collagen area to the total tissue area and expressed as a percentage^14^.

### Prediction of Transcription Factor Binding Sites

To identify potential upstream transcriptional regulators of the downstream gene, we performed transcription factor binding site prediction analysis using the JASPAR database (https://jaspar.genereg.net/). First, the reference sequence of the gene’s promoter region, spanning from 2000 bp upstream to 100 bp downstream of the transcription start site, was obtained via the UCGC Genome Browser. This sequence was submitted to the JASPAR online tool for analysis. To screen for high-confidence binding sites, the Absolute Score Threshold was set to 700. All predicted binding sites output by the system were required to meet this threshold. Subsequently, the corresponding candidate transcription factors were ranked according to the scores of their binding sites, and further screened in combination with their known biological functions.

### Chromatin immunoprecipitation (ChIP)

ChIP assay was performed using a commercial ChIP Assay Kit (Beyotime, Shanghai, China; Cat# P2078) according to the manufacturer’s instructions. Briefly, cells were cross-linked with 1% formaldehyde for 10 min at room temperature, and the reaction was quenched with 125 mM glycine. Cells were then harvested, lysed, and sonicated to shear chromatin into fragments of 100–1000 bp, as confirmed by agarose gel electrophoresis. After centrifugation, the supernatant containing fragmented chromatin was diluted and pre-cleared with protein A/G agarose beads. An aliquot of the diluted chromatin was saved as input control. The remaining chromatin was incubated overnight at 4 °C with 2 µg of anti-NR5A1 antibody (Proteintech, Wuhan, China) or normal rabbit IgG (negative control). Immune complexes were captured by adding protein A/G agarose beads and incubated for 2 h at 4 °C with rotation. Beads were washed sequentially with low-salt, high-salt, and LiCl wash buffers, followed by TE buffer, as provided in the kit. Bound chromatin was eluted and reverse cross-linked by heating at 65 °C overnight. DNA was purified using the DNA purification kit provided. Enrichment of the PPARγ promoter region was analyzed by PCR using primers. PCR products were resolved on 2% agarose gels and visualized by ethidium bromide staining. Band intensities were quantified by densitometry using ImageJ software, and the percentage of input was calculated as (ChIP signal / input signal) × 100%.

### Measurement of serum lipid profiles

Blood samples were collected from mice via the retro-orbital sinus at 1 week and 4 weeks after the initiation of BAPN treatment (i.e., at 5 and 8 weeks of age). Four groups of mice (n=15 per group) were included at each time point. Blood was allowed to clot at room temperature for 30 min, and serum was separated by centrifugation at 3,000 × g for 15 min at 4 °C. Serum samples were stored at −80 °C until analysis. Serum levels of total cholesterol (T-CHO), triglycerides (TG), high-density lipoprotein cholesterol (HDL-C), and low-density lipoprotein cholesterol (LDL-C) were measured using a fully automated biochemical analyzer with commercially available assay kits according to the manufacturer’s protocols. All measurements were performed by Servicebio (Wuhan, China).

### Statistical analysis

Data analysis was performed using R (version 4.3.2) and GraphPad Prism (version 9.0) software. The raw proteomic and transcriptomic sequencing data of plasma samples were obtained from our previously published study^15^. In the present study, the raw data were re-analyzed using R to evaluate the differential expression of APOH. Continuous variables are presented as mean ± SD or as box-and-whisker plots (indicating the 75th and 25th percentiles where applicable). Categorical variables are expressed as numbers and percentages. Group comparisons were performed using two-sided t-test, Mann-Whitney U test, or chi-square test, as appropriate. For multiple group comparisons, one-way ANOVA followed by Tukey’s or Dunnett’s post hoc test was applied; alternatively, the Kruskal-Wallis test followed by Dunn’s multiple comparisons test was conducted when assumptions were violated. For time-to-event data, survival analysis was performed using the Kaplan-Meier method, and differences between groups were compared by the log-rank test. Statistical significance was set at P < 0.05.

## RESULTS

### APOH is elevated in circulation and accumulates in aortic tissues in AD

To identify key circulating factors involved in the initiation and progression of AD, we previously performed integrated transcriptomic and proteomic analyses of plasma samples from patients with AD and healthy controls. The results revealed significantly differentially expressed genes between AD patients and controls, with APOH ranking among the top 10 most significantly altered genes (Fig. 1A). Proteomic analysis also identified multiple differentially expressed proteins (Fig. 1B). Although APOH did not rank within the top 10 in the proteomic dataset, it showed an overall upregulated trend, which was further validated by heatmap analysis (Fig. 1C). Venn diagram analysis showed that a total of 32 genes/proteins were consistently altered at both the mRNA and protein levels, including 26 upregulated and 6 downregulated (Fig. 1D). Both the mRNA and protein levels of APOH were upregulated. These results suggest that APOH may serve as a circulating regulatory factor associated with AD. To validate the clinical relevance of APOH, we further measured its plasma levels. ELISA results showed that plasma APOH concentrations were significantly elevated in AD patients (Fig. 1E). However, APOH mRNA expression in aortic tissues did not differ significantly between the AD group and the normal control group (Fig. 1F), suggesting that APOH is not primarily derived from local vascular synthesis but rather from hepatic synthesis, as previously reported^16^. Although mRNA levels remained unchanged, western blot analysis revealed a significant increase in APOH protein expression in AD aortic tissues (Fig. 1G), indicating possible post-transcriptional regulation or deposition from the circulation. Histological analysis further confirmed marked structural damage in AD tissues, including disruption of vascular architecture (HE staining), elastic fiber disruption (EVG staining), and increased collagen deposition (Masson’s trichrome staining) (Fig. 1H). Moreover, IHC and IF analyses showed significant enrichment of APOH in AD aortic tissues, with both positive expression and fluorescence intensity being markedly higher than those in the control group (Fig. 1H). Taken together, APOH is significantly elevated in the circulation and deposited in aortic tissues during AD, suggesting that it acts as a blood-derived regulatory factor involved in the pathogenesis and progression of the disease.

**Fig. 1.**
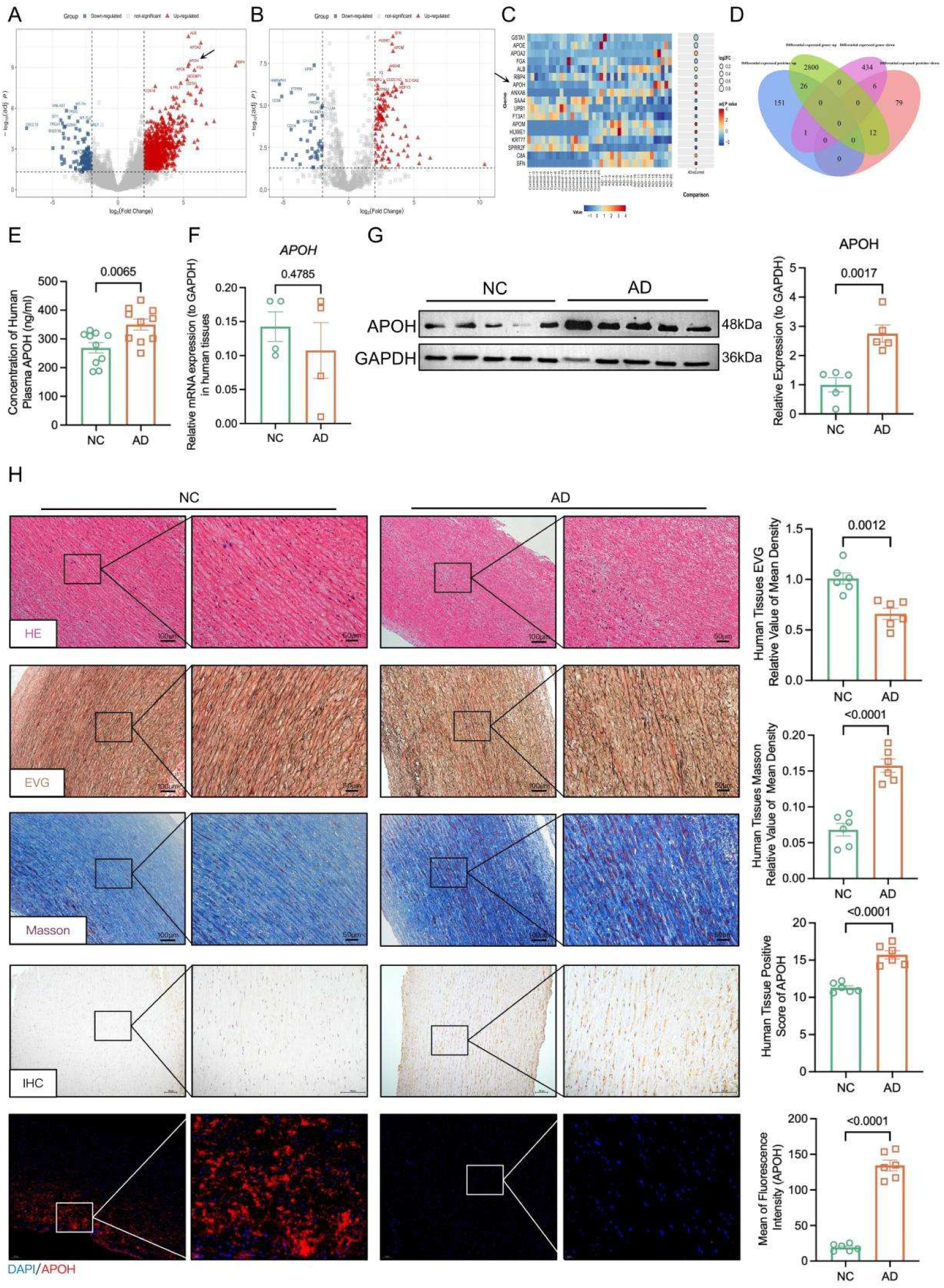
APOH is upregulated in human aortic dissection with vascular wall remodeling. (A, B) Volcano plots of differentially expressed genes and proteins from plasma exosome RNA-seq and proteomics, respectively; red dots represent upregulated genes/proteins, blue dots represent downregulated genes/proteins, with TOP10 differentially expressed genes/proteins labeled. (C) Heatmap of *APOH* and other differentially expressed genes. (D) Venn diagram illustrating the concordant changes between transcriptomic and proteomic data. (E) Plasma APOH levels in AD patients versus normal controls (ELISA) (*n* = 10). (F) *APOH* mRNA expression in AD versus normal control aortic tissues (qRT-PCR) (*n* = 4). (G) APOH protein expression in aortic tissues (WB) (*n* = 5). (H) Representative images of H&E staining, EVG staining, Masson’s trichrome staining, and APOH IHC and IF of aortic tissues (*n* = 6).

### APOH knockdown alleviates BAPN/Ang-II–induced AD and vascular remodeling in vivo

To investigate the functional role of APOH in AD, we established a BAPN/Ang-II - induced mouse model of AD and performed AAV-mediated knockdown of APOH (Fig. 2A). We first validated the efficiency of APOH knockdown. Liver qPCR analysis showed that APOH mRNA levels were increased in the AD model group, and administration of AAV-APOH reduced APOH mRNA levels (Fig. 2B). Plasma measurements also confirmed a decrease in circulating APOH levels (Fig. 2C). KM survival analysis showed that APOH knockdown significantly improved the survival rate of AD model mice (Fig. 2D). Gross morphological examination revealed that APOH knockdown reduced the severity and incidence of AD, which was further confirmed by quantitative analysis (Fig. 2E-F). In mice treated with BAPN, approximately 40% developed ascending aortic aneurysms, 20% developed descending aortic aneurysms, and some exhibited both types. Echocardiography primarily measures the internal diameter of the ascending aorta. Echocardiography demonstrated that APOH knockdown significantly attenuated BAPN/Ang-II-induced aortic dilation, as reflected by a decreased ascending aortic diameter (Fig. 2G). Serum lipid profile analysis indicated that BAPN treatment induced marked metabolic disturbances, characterized by elevated levels of total cholesterol (TC) and low-density lipoprotein (LDL). APOH knockdown partially ameliorated these changes, particularly at the 4-week time point, although levels did not return to normal (Fig. 2H). Histological analysis further showed that APOH knockdown significantly improved vascular structural damage. HE staining revealed more intact vascular wall architecture, EVG staining showed reduced elastic fiber disruption, and Masson’ s trichrome staining indicated decreased collagen deposition, suggesting alleviation of extracellular matrix remodeling (Fig. 2I). Immunofluorescence results further demonstrated marked deposition of APOH in the aortic wall of AD mice, which was significantly reduced by APOH knockdown. Of note, APOH exhibited clear colocalization with the vascular smooth muscle cell marker SM22, suggesting that circulating APOH may exert effects on vascular smooth muscle cells. This colocalization signal was markedly attenuated following APOH knockdown, accompanied by partial restoration of vascular structure (Fig. 2I). Collectively, these findings indicate that APOH promotes AD development *in vivo*, and its effect may be mediated, at least in part, through direct action on vascular smooth muscle cells, rather than solely via systemic metabolic regulation.

**Fig. 2.**
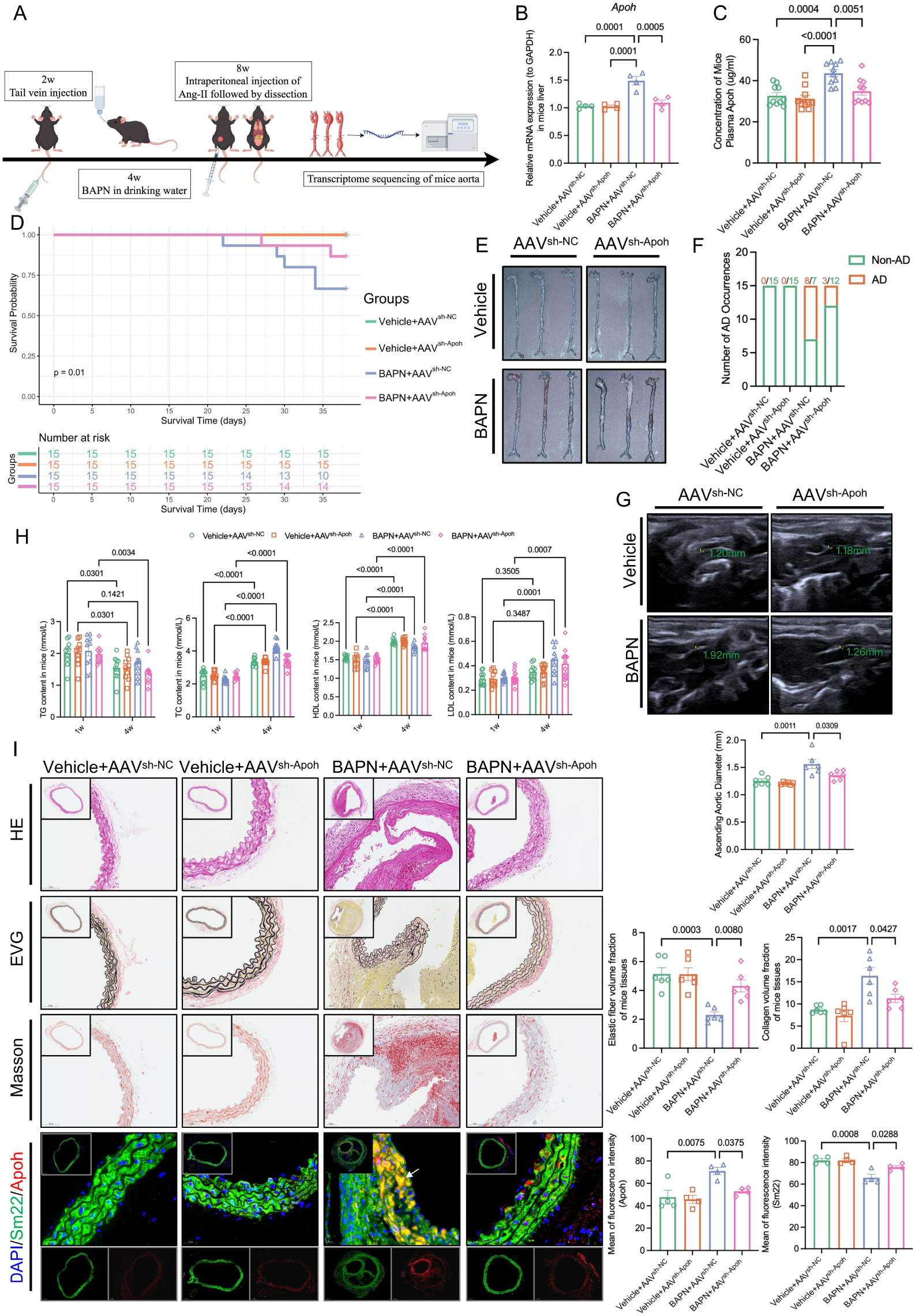
APOH knockdown attenuates Ang-II-induced aortic dissection and ameliorates vascular wall remodeling in mice. (A) Schematic illustration of the experimental protocol. (B) qPCR results demonstrating reduced expression efficiency in mouse liver (n = 4). (C) Apoh levels in circulating plasma of mice (n = 10). (D) KM survival curves; number at risk at each time point is shown below (n = 15). (E) Representative gross images of ascending aortas. (F) Incidence of AD across experimental groups. (G) Representative B-mode ultrasound images of ascending aortic diameter (mm) with measurement regions indicated in green, and corresponding quantitative analysis (n = 6). (H) Dynamic changes in serum TG, TC, HDL-C, and LDL-C levels at 1 and 4 weeks (n = 10). (I) Representative images of H&E, EVG, and Masson’ s trichrome staining, and DAPI/Sm22/Apoh triple immunofluorescence of aortic tissues; right panels: quantification of elastic fiber, collagen, and Apoh fluorescence intensity (n = 6).

### Transcriptomic profiling identifies PPAR signaling as a key pathway mediating APOH-associated vascular remodeling

To elucidate the molecular mechanism underlying APOH-mediated vascular remodeling, we first examined the expression of VSMC phenotype markers in aortic tissues. BAPN treatment significantly induced phenotypic switching, characterized by decreased expression of contractile markers (ACTA2 and TAGLN) and increased expression of the synthetic marker OPN. Notably, APOH knockdown partially reversed these changes, restoring contractile marker expression and reducing OPN levels (Fig. 3A), suggesting that APOH is involved in regulating VSMC phenotypic switching. To further explore the underlying molecular mechanisms, we performed transcriptome sequencing of aortic tissues from BAPN-treated mice (AAV^sh-NC^ vs. AAV^sh-Apoh^). Differential expression analysis revealed extensive transcriptional alterations following APOH knockdown (Fig. 3B). GO functional enrichment analysis showed that the differentially expressed genes were mainly enriched in mitochondrial function, redox processes, and cytoskeletal/extracellular matrix regulation (Fig. 3C). KEGG pathway analysis further indicated that these genes were primarily enriched in metabolic and cardiovascular-related pathways, including oxidative phosphorylation, fatty acid metabolism, vascular smooth muscle contraction, and ECM–receptor interaction (Fig. 3D). Overall, this enrichment pattern was not dominated by a single pathway but rather reflected a coordinated reprogramming of metabolic regulation and contractile function, which is typically finely tuned by nuclear receptor signaling pathways. Notably, the PPAR signaling pathway was also identified among the enriched terms, suggesting that lipid metabolism-related transcriptional regulation may be involved in APOH-mediated vascular remodeling. Among the nuclear receptor pathways, PPARγ (encoded by PPARG) is a master regulator of vascular smooth muscle cell phenotype switching, lipid metabolism, and inflammation, and its downstream targets were prominently enriched in our dataset. Therefore, we selected PPARG as the candidate for further mechanistic investigation. To further investigate the upstream regulatory mechanism of PPARγ, transcription factor binding site prediction was performed using the JASPAR database. The results revealed that multiple transcription factors exhibited high binding scores within the PPARγ promoter region, among which NR5A1 showed high-confidence predicted binding sites (Fig. 3F), suggesting that NR5A1 may act as an upstream transcriptional regulator of PPARγ. Meanwhile, protein-protein interaction network analysis revealed potential associations among APOH, PPARγ, NR5A1, and VSMC phenotype markers (Fig. 3G), supporting the existence of a coordinated regulatory network. Collectively, these findings suggest that APOH may participate in VSMC phenotypic switching and vascular remodeling through the regulation of PPAR-related transcriptional programs.

**Fig. 3.**
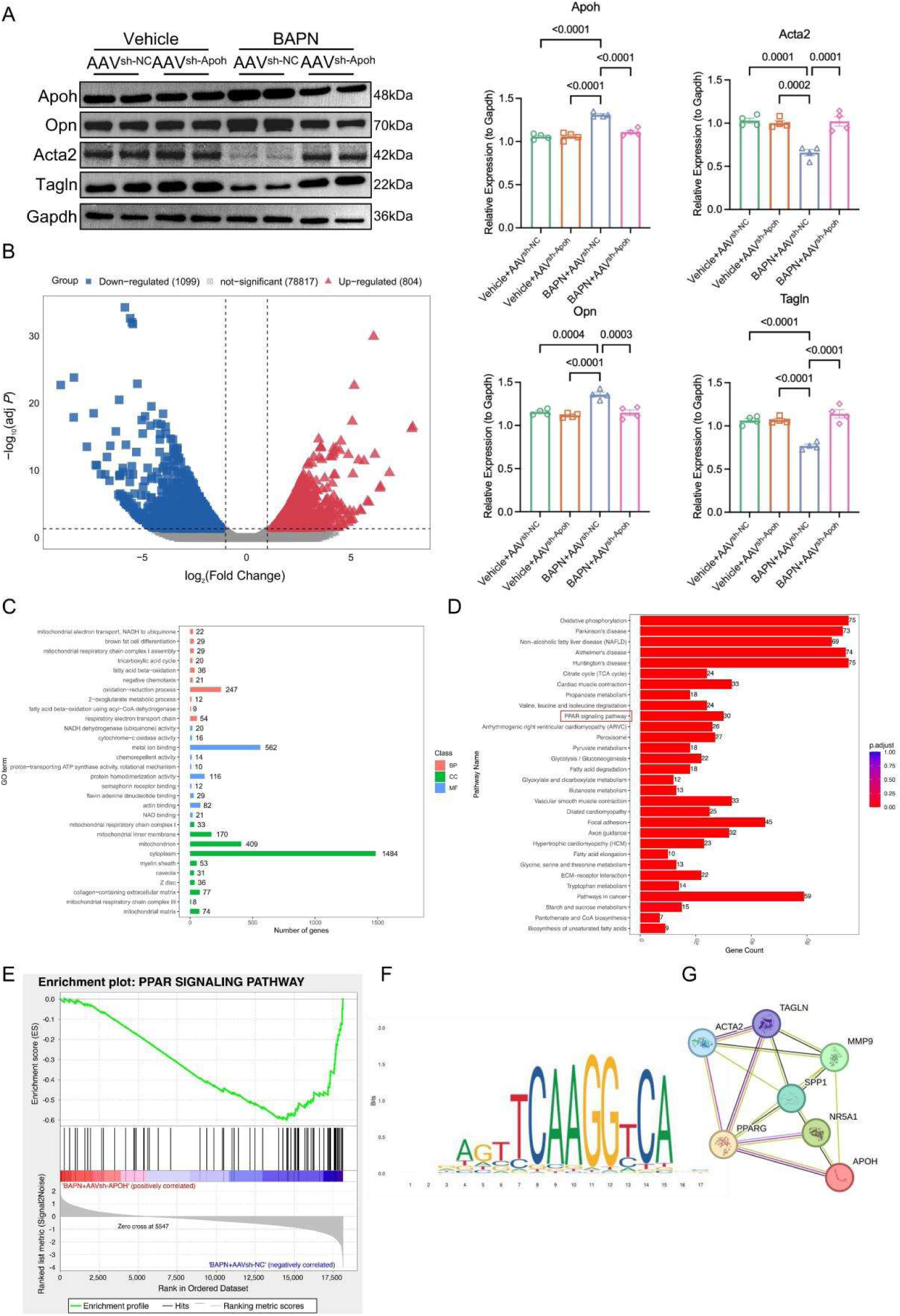
APOH knockdown modulates vascular smooth muscle cell phenotypic switching through PPAR signaling pathway. (A) WB validation of AAV^sh-Apoh^ knockdown efficiency and its effects on VSMC phenotypic markers Opn, Acta2, and Tagln (n = 4). (B) Transcriptome sequencing results from mouse aortic tissues (n = 5), volcano plot of differentially expressed genes: blue dots represent downregulated genes (1,099), gray dots represent non-significant genes (78,817), red dots represent upregulated genes (804). (C) Representative results of GO enrichment analysis: significantly enriched terms in biological process (BP), cellular component (CC), and molecular function (MF). (D) Representative results of KEGG pathway enrichment analysis; red indicates high significance. (E) Enrichment analysis of PPAR signaling pathway: green line represents enrichment curve; bar plot shows associated genes. (F) Sequence analysis of NR5A1 binding site in PPARγ promoter region. (G) Protein-protein interaction network of APOH with PPARγ, NR5A1, and VSMC phenotypic markers.

### APOH promotes HAVSMC phenotypic switching via the NR5A1–PPARγ axis

*In vivo* experiments, immunohistochemical staining revealed that BAPN treatment significantly upregulated the expression of NR5A1, PPARγ, and its downstream lipid metabolism-related protein FABP4 in the aorta compared with the Vehicle group (Fig. 4A). Notably, APOH knockdown markedly decreased the expression levels of all the above proteins, suggesting that APOH positively regulates the NR5A1–PPARγ signaling axis *in vivo*. To investigate the direct effect of APOH on HAVSMC, we stimulated cells with Ang-II and recombinant APOH protein. The results showed that Ang-II significantly induced a phenotypic switch of HAVSMC from a contractile to a synthetic phenotype, as evidenced by decreased expression of the contractile markers ACTA2 and TAGLN, and increased expression of the synthetic and inflammation-related markers OPN and MMP9 (Fig. 4B). Of note, APOH treatment alone induced similar changes and even showed a stronger trend for some markers, suggesting that APOH can mimic or even enhance Ang-II-induced phenotypic switching. To further elucidate the molecular mechanism underlying this process, we knocked down NR5A1 or PPARγ in the presence of APOH stimulation. The results showed that siNR5A1 or siPPARγ each significantly reversed APOH-induced phenotypic switching: the expression of ACTA2 and TAGLN was restored, whereas the expression of OPN, MMP9, and FABP4 was markedly reduced (Fig. 4C). These findings indicate that NR5A1 and PPARγ play critical roles in APOH-mediated phenotypic switching in HAVSMC. Furthermore, ChIP assay results showed that NR5A1 directly binds to the PPARγ promoter region, and this binding was significantly enhanced upon Ang-II stimulation (Fig. 4D), confirming that NR5A1 acts as an upstream regulator of PPARγ. To determine the pro-inflammatory effect of APOH in aortic dissection, we examined the expression of key inflammatory factors in the aortic tissues of AD model mice. As shown in Fig. 5A, compared with the control group, the BAPN+AAV^sh-NC^ group exhibited significantly elevated mRNA expression of the pro-inflammatory cytokines TNF-α, IL-6, and MCP1 in the aorta, whereas APOH knockdown effectively blocked BAPN/Ang-II-induced aortic inflammation. In the *in vitro* HAVSMC model, treatment with recombinant APOH significantly upregulated the mRNA expression of TNF-α, IL-6, and MCP1 compared with the Con group, while siNR5A1 or siPPARγ significantly suppressed APOH-induced inflammatory responses. Collectively, these results demonstrate that in the pathological context of AD, APOH promotes HAVSMC phenotypic switching and inflammation via the NR5A1-PPARγ axis, and blockade of this axis effectively inhibits these pathological processes.

**Fig. 4.**
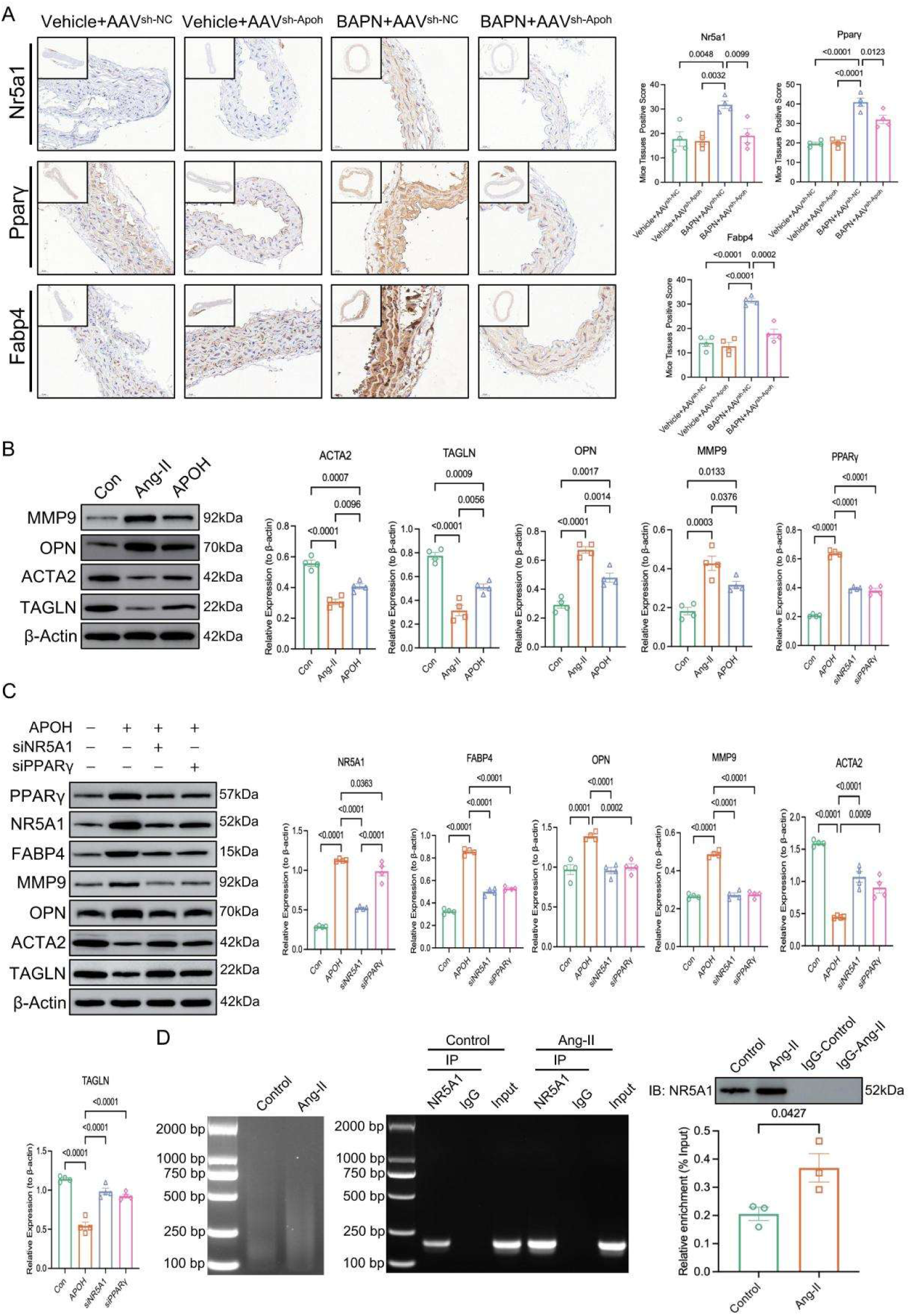
APOH-NR5A1-PPARγ axis regulates vascular smooth muscle cell phenotypic switching. (A) IHC staining showing expression of Nr5a1, Ppar γ, and Fabp4 in aortic tissues from different groups (n = 4); quantitative analysis shown on the right. (B) WB and quantitative analysis: Effects of Ang-II and APOH treatment on expression of VSMC phenotypic markers (ACTA2, TAGLN, OPN, MMP9) (n = 4). (C) WB and quantitative analysis: Regulatory effects of APOH on downstream proteins and VSMC phenotypic markers after knockdown of NR5A1 or PPARγ (n = 4). (D) ChIP validation of direct binding between NR5A1 and PPARγ promoter: sonication efficiency, PCR electrophoresis, and relative enrichment analysis (n = 3).

**Fig. 5.**
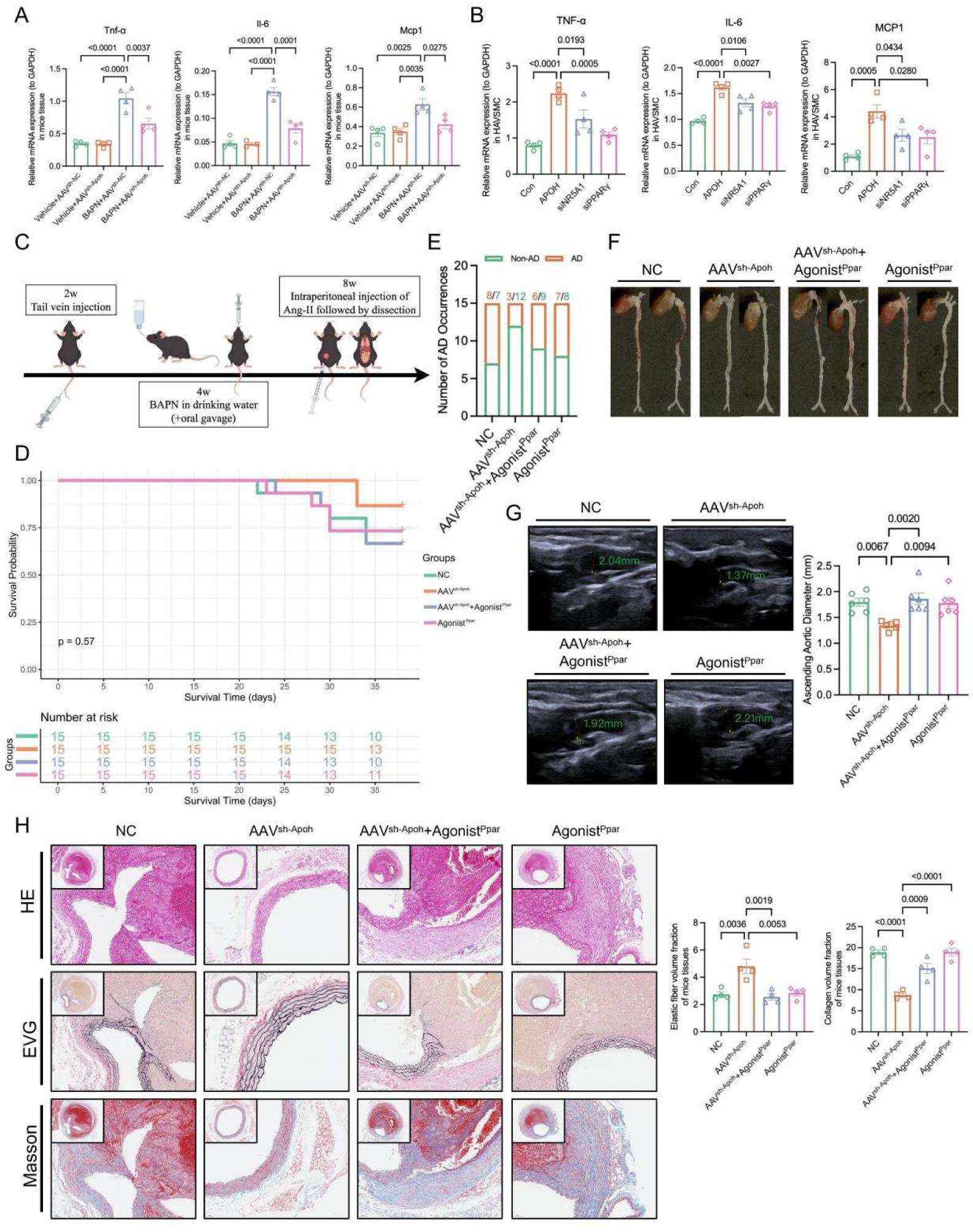
PPARγ agonist abrogates the protective effect of APOH deficiency against AD in mice. (A) Quantitative PCR analysis of inflammatory cytokine (Tnf-α, Il-6, Mcp1) mRNA expression levels in aortic tissues from different experimental groups (n = 4). (B) qPCR validation of pro-inflammatory mediators (TNF-α, IL-6, MCP1) expression in HAVSMC under APOH treatment and NR5A1/PPARγ knockdown conditions (n = 4). (C) Schematic illustration of the experimental design protocol. (D) KM survival curve analysis; number of animals at risk at each time point shown below the graph (n = 15). (E) Incidence rates of AD across experimental groups. (F) Representative gross anatomical images of ascending aortas from different groups. (G) Representative B-mode ultrasound images of ascending aortic diameter (mm) with measurement regions indicated by green boxes, and corresponding quantitative analysis (n = 6). (H) Representative images of aortic cross-sections stained with H&E, EVG for elastic fibers, and Masson’s trichrome; right panels show quantitative analysis of elastic fiber integrity and collagen deposition (n = 4).

### PPARγ activation functionally reverses the protective effects of APOH silencing in vivo

To functionally validate the role of PPARγ in APOH-mediated AD pathogenesis, we performed a rescue experiment by administering a PPARγ agonist to BAPN-treated mice with APOH knockdown (Fig. 5C). KM survival analysis showed no statistically significant difference in survival rates among the groups (p=0.57) (Fig. 5D). However, APOH silencing significantly reduced the incidence of aortic dissection, whereas PPARγ agonist treatment markedly attenuated this protective effect (Fig. 5E–F). Echocardiography results demonstrated that APOH silencing significantly inhibited ascending aortic dilation, while PPARγ activation partially reversed the reduction in aortic diameter (Fig. 5G). Histological analysis revealed that APOH silencing substantially ameliorated aortic structural damage, as evidenced by reduced elastic fiber disruption and decreased collagen deposition; in contrast, PPARγ activation weakened this protective effect, leading to aggravated elastic fiber destruction and re-accumulation of collagen (Fig. 5H). These findings provide functional evidence that PPARγ mediates the pro-pathogenic role of APOH in AD *in vivo*.

## DISCUSSION

Through multi-dimensional experimental evidence, this study systematically elucidates the key role of APOH in the pathogenesis of AD and its underlying molecular mechanisms. We found that APOH is elevated in the circulation and accumulates in aortic tissues, and that knockdown of APOH ameliorates the AD phenotype. Transcriptomic analysis identified the PPAR signaling pathway as a key mediator. Further investigation revealed that APOH promotes vascular smooth muscle cell phenotypic switching and inflammatory responses via the NR5A1–PPARγ axis, and functional rescue experiments confirmed that PPARγ mediates the pro-pathogenic effect of APOH. These findings provide new insights into the pathogenesis of AD and the development of targeted therapeutic strategies.

The present study reveals a dual feature of APOH in AD patients: elevated circulating levels and tissue accumulation. Previous studies have shown that APOH (also known as β2-glycoprotein I) is a plasma protein primarily synthesized by the liver and involved in various physiological processes, including coagulation regulation, lipid metabolism, and inflammatory responses^17^. In autoimmune diseases such as antiphospholipid syndrome, APOH acts as an autoantigen^18^. However, the role of APOH in cardiovascular diseases, particularly AD, has not yet been reported.

Our study found that although APOH mRNA expression in aortic tissues did not change significantly, the protein level was markedly elevated, suggesting that circulating APOH may accumulate in the aortic wall through extracellular deposition. This finding echoes previous studies on the role of APOH in atherosclerosis, which demonstrated that APOH promotes foam cell formation by binding to oxidized low-density lipoprotein^19,20^. Tabuchi et al. further showed that the C-reactive protein/oxidized low-density lipoprotein/β2-glycoprotein I complex induces lipid accumulation and inflammatory responses in macrophages via the p38/MAPK and NF-κB signaling pathways, providing mechanistic evidence for the role of APOH in vascular inflammation^21^. Nevertheless, our study reveals a pathogenic role of APOH in acute aortic disease, extending the understanding of the biological functions of APOH.

Of note, the colocalization of APOH with the VSMC marker SM22 suggests that circulating APOH may directly act on vascular smooth muscle cells. This finding provides new clues for understanding how circulating factors regulate vascular cell function. Previous studies have primarily focused on the role of locally synthesized cytokines and growth factors in vascular diseases, whereas our study highlights the direct regulatory role of circulating proteins in acute vascular injury.

This study reveals the pro-inflammatory properties of the NR5A1–PPARγ axis in the pathological context of AD. Traditionally, PPARγ, as a member of the nuclear receptor superfamily, is recognized to exert anti-inflammatory effects in metabolic diseases and inflammatory responses^22^. However, our study found that in the acute injury microenvironment of AD, PPARγ switches to a pro-inflammatory effector, challenging conventional understanding and revealing the context-dependent nature of nuclear receptor functions. Transcriptomic analysis showed that APOH knockdown significantly altered the expression of genes related to mitochondrial function, redox balance, and the cytoskeleton, suggesting that APOH influences VSMC function by regulating cellular metabolism and structure. Gene set enrichment analysis revealed significant enrichment of the PPAR signaling pathway. Combined with JASPAR prediction and ChIP validation of direct binding of NR5A1 to the PPARγ promoter, this study establishes the APOH–NR5A1–PPARγ signaling axis, providing a new perspective for understanding the mechanisms of vascular remodeling. This may explain why PPARγ exhibits pro-inflammatory properties in AD, possibly due to aberrant activation of its upstream regulator NR5A1.

VSMC phenotypic switching is a key process in vascular remodeling, and the transition from a contractile to a synthetic phenotype leads to extracellular matrix degradation, release of inflammatory cytokines, and disruption of vascular structure ^23^. Our study demonstrated that APOH promotes VSMC phenotypic switching via the NR5A1– PPARγ signaling axis, as evidenced by the downregulation of contractile markers and upregulation of synthetic markers. Of note, the role of PPARγ in vascular diseases varies significantly: in atherosclerosis, a chronic inflammatory process, PPARγ agonists exhibit anti-inflammatory and anti-proliferative effects^24^. However, in the context of acute injury in AD, our study found that PPARγ activation conversely promotes disease progression. This functional discrepancy may be attributed to distinct pathological microenvironments—chronic inflammation versus acute vascular injury—leading to functional reprogramming of nuclear receptor signaling pathways, thereby resulting in markedly different cellular responses to the same signal. Furthermore, our study revealed that APOH-induced inflammatory responses are closely associated with phenotypic switching, suggesting that inflammation and phenotypic switching may synergistically promote AD progression. This finding is consistent with recent studies on the role of vascular inflammation in AD^25^.

Despite the important findings of this study, several limitations should be acknowledged. First, our study is primarily based on a mouse model. Although mouse models are widely used in vascular disease research, differences between murine models and human diseases must be interpreted with caution. Future studies are warranted to validate our findings in larger clinical cohorts. Second, although we demonstrated that NR5A1 binds to the PPARγ promoter, the precise mechanism by which APOH activates NR5A1 remains unclear. Further studies are needed to explore the receptor of APOH and its downstream signaling pathways. Third, our study mainly focused on the role of the NR5A1–PPARγ axis in VSMC; however, the function of this axis in other cell types, such as endothelial cells and macrophages, requires further investigation.

## CONCLUSION

This study systematically elucidates the molecular mechanism by which APOH promotes the development and progression of AD through activation of the NR5A1– PPARγ axis. Our findings not only provide new insights into the pathogenesis of AD but also lay a theoretical foundation for the development of targeted therapeutic strategies. These results highlight the direct regulatory role of circulating factors in acute vascular diseases, as well as the context-dependent nature of nuclear receptor signaling pathway functions, which have important implications for vascular biology and the treatment of cardiovascular diseases.

## Glossary

AD: aortic dissection
ECM: extracellular matrix
HAVSMC: human aortic vascular smooth muscle cell
VSMC: vascular smooth muscle cell
HDL: high-density lipoprotein
HDL-C: high-density lipoprotein cholesterol
LDL: low-density lipoprotein
LDL-C: low-density lipoprotein cholesterol
TC: total cholesterol
T-CHO: total cholesterol
TG: triglycerides

## Acknowledgements

No.

## Author contributions

Likang Ma and Zhihuang Qiu designed and supervised the study. Likang Ma and Lei Jin performed formal analysis, investigation and drafted the manuscript. Likang Ma and Jiakang Li contributed to the methodology and the visualization. Likang Ma and Lei Jin performed the *in vivo* experiments. Lei Jin and Juncheng Liu performed the *in vitro* experiments. Lei Jin performed the AAV administration. Maolin Liu performed the informatics analysis. Liangwan Chen and Zhihuang Qiu contributed to the writing, review, and editing the manuscript. All authors contributed to the article and approved the submitted version.

## Ethics declaration

The human research protocol was approved by the Ethics Committee of Fujian Medical University Union Hospital [No. 2023-96]. Animal experiments were approved by the Animal Ethics Committee of Fujian Medical University [IACUC FJMU 2023-Y-0269] and complied with the ARRIVE guidelines.

